# Fgf8 promotes survival of nephron progenitors by regulating BAX/BAK-mediated apoptosis

**DOI:** 10.1101/2022.08.25.505319

**Authors:** Matthew J. Anderson, Salvia Misaghian, Nirmala Sharma, Alan O. Perantoni, Mark Lewandoski

**Affiliations:** Genetics of Vertebrate Development Section, Cancer and Developmental Biology Laboratory, National Cancer Institute, National Institutes of Health, Frederick, MD 21702, USA; Renal Differentiation and Neoplasia Section, Cancer and Developmental Biology Laboratory, National Cancer Institute, National Institutes of Health, Frederick, MD 21702, USA

**Keywords:** Fibroblast growth factor, Fgf8, Kidney, Apoptosis, Cell Death

## Abstract

Fibroblast growth factors (*Fgfs*) have long been implicated in processes critical to embryonic development, such as cell survival, migration, and differentiation. Several mouse models of organ development ascribe a prosurvival requirement specifically to FGF8. Here, we explore the potential role of prosurvival FGF8 signaling in kidney development. We have previously demonstrated that conditional deletion of *Fgf8* in the mesodermal progenitors that give rise to the kidney leads to renal aplasia in the mutant neonate. Deleterious consequences caused by loss of FGF8 begin to manifest by E14.5 when massive aberrant cell death occurs in the cortical nephrogenic zone in the rudimentary kidney as well as in the renal vesicles that give rise to the nephrons. To rescue cell death in the *Fgf8* mutant kidney, we inactivate the genes encoding the pro-apoptotic factors BAK and BAX. In a wild-type background, the loss of *Bak* and *Bax* abrogates normal cell death and has minimal effect on renal development. However, in *Fgf8* mutants, the combined loss of *Bak* and *Bax* rescues aberrant cell death in the kidneys and restores some measure of kidney development: 1) the nephron progenitor population is greatly increased; 2) some glomeruli form, which are rarely observed in *Fgf8* mutants; and 3) kidney size is rescued by about 50% at E18.5. The development of functional nephrons, however, is not rescued. Thus, FGF8 signaling is required for nephron progenitor survival by regulating BAK / BAX and for subsequent steps involving, as yet, undefined roles in kidney development.

## Introduction

Fibroblast growth factor (FGF) signaling is essential for the development of numerous tissues during embryonic development (reviewed [1]). In some tissues, a deficiency of FGF signaling causes aberrant cell death and subsequent tissue loss or malformation [2–6]. We have previously shown that conditional inactivation of *Fgf8* in the nascent mesoderm using TCre (in TCre; *Fgf8^flox/null^* embryos; see Material and Methods for genetic cross), which deletes *Fgf8* before it is expressed in the kidney primordium, results in neonatal lethality as a result of kidney aplasia [6]. This failure of kidney development appears to stem from some combination of extensive cortical and tubular cell death and a failure of *Lhx1* expression. *Lhx1* encodes a Lim-homeobox transcription factor that is expressed in the ureteric bud, nephronic pretubular aggregates and subsequent renal vesicles [7]. Conditional deletion of *Lhx1* in the metanephric mesenchyme lineage results in loss of nephron structures including glomeruli and associated nephron tubules; however, NPs appear to be unaffected [7, 8]. In TCre; *Fgf8^flox/null^* mutants, the renal vesicles fail to express *Lhx1* [6]. Another feature of the TCre; *Fgf8^flox/null^* mutant kidney is widespread *c*ell death in the cortical region where the nephron progenitors (NPs) reside as well as in the newly formed renal vesicles beginning around embryonic day 14 (E14). Therefore, *Fgf8* may be responsible for two separable actions: 1) maintaining cell survival of the NPs and 2) initiation of *Lhx1* expression in the forming renal vesicles.

Cell death occurs through numerous mechanisms, which are used for classifying this phenomenon into three main types: apoptosis, necrosis, and autophagy (reviewed [9]). Apoptosis involves an intercellular cascade that ultimately leads to fragmentation of the DNA. This cascade can be initiated intrinsically or extrinsically. The extrinsic pathway is initiated by “death ligands,” such as TNFα, binding to cell surface receptors to trigger death of the cell (reviewed in [9]). The intrinsic pathway is initiated by a cytotoxic stimulus, such as withdrawal of a trophic signaling molecule, which initiates a cascade that first impacts integrity of the mitochondria. The Bcl-2 family of proteins are the major factors which interact to initiate or suppress cell death by this mitochondrial-mediated intrinsic pathway [10]. Tilting the balance of expression and activation of pro-death versus pro-survival Bcl-2 family members results in mitochondrial pore formation and subsequent cell death. When upstream interactions induce a cell death response, BAK and BAX oligomerize and form pores in the mitochondria. Apoptosis still occurs when either *Bak* or *Bax* are individually deleted; however, deletion of both genes blocks apoptosis [11, 12]; thus, *Bak* and *Bax* are redundantly required for the intrinsic apoptotic pathway. To study the effects of cell death during development due to specific genetic alterations, such as deletions of *Fgf3, Kif20b, Sirt6*, or *Tbx1*, cell survival has been genetically restored by the deletion of genes encoding proapoptotic factors, such as *Bak* and *Bax* or *p53* [2, 13–15]. In the current study, we examine the function of FGF8 signaling during embryonic kidney development and resolve its relative roles in cell survival and subsequent kidney development by inactivating the proapoptotic genes, *Bak* and *Bax*.

## Results

### *Fgf8* is required for survival of metanephric mesenchyme

We have previously shown that kidneys in which *Fgf8* is genetically removed show extensive cell death within the cortical nephrogenic zone beginning around E14.5 [6]. This zone includes the nephron progenitor population marked by the homeobox gene *Six2* [16]. To determine if cell death in TCre; *Fgf8^flox/null^* mutant kidneys occurs in this critical population, we evaluated the SIX2-expressing cells by immunofluorescence staining and the dying cells with Lysotracker red [17]. As expected, there is very little cell death in the cortical regions of E14.5 control kidneys (Figure 1A-C); however, mutants display extensive cell death in this region (Figure 1D-F). To quantify co-localized SIX2 and Lysotracker staining, spatial models were built and analyzed using Imaris software (see Materials and Methods). These analyses reveal that, on average, 13.4% of SIX2-positive cells are dying in *Fgf8* mutants at E14.5, compared to less than 2% of SIX2-positive cells in controls (Figure 1G). Additionally, 84.7% of dying cells within the cortex are SIX2 positive in TCre; *Fgf8^flox/null^*. Therefore, at E14.5, when cell death is beginning in *Fgf8* mutants, a substantial number of nephron progenitors (NPs) are dying and a majority of the aberrant cell death that is observed in the *Fgf8* mutant is occurring in the SIX2-positive NPs.

**Figure 1.**
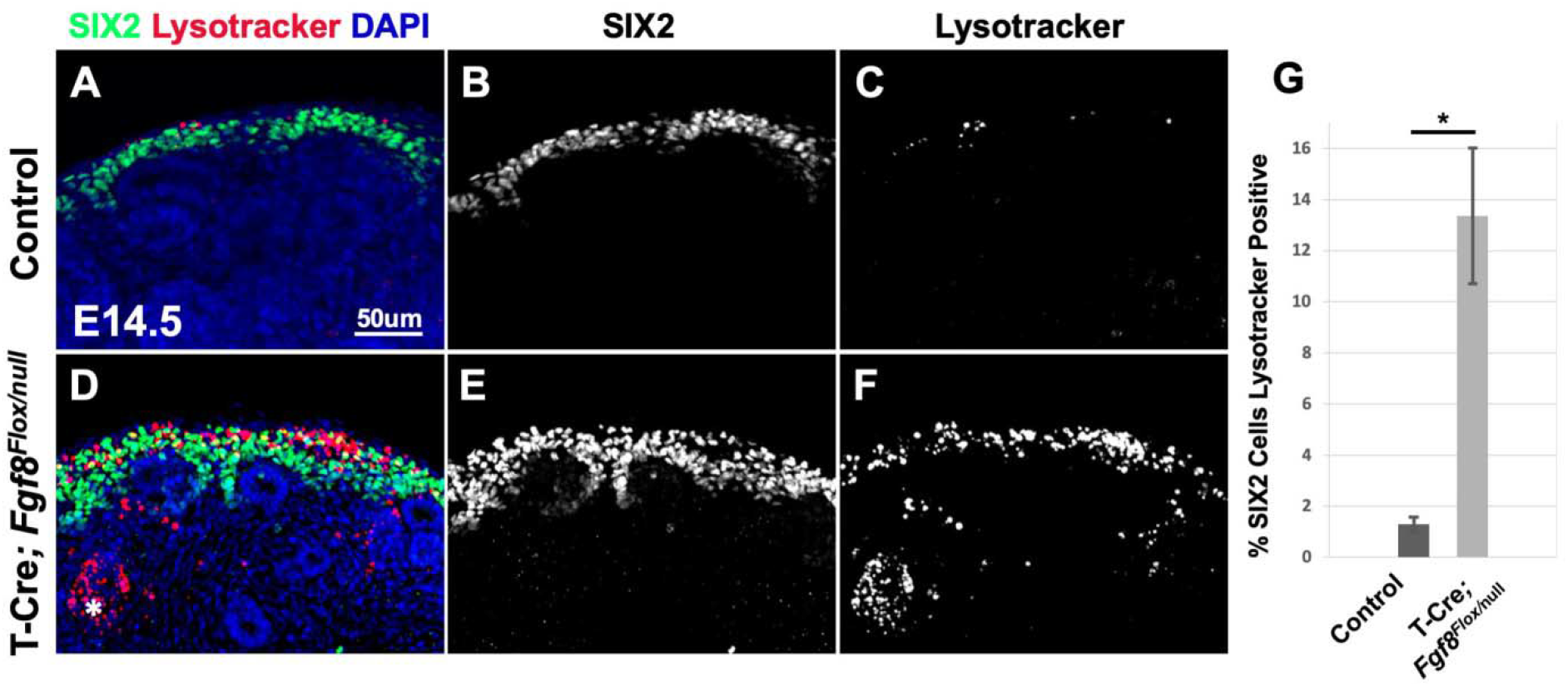
Loss of *Fgf8* causes cell death in nephron progenitors. **A-F)** Max intensity projection of mid-sagittal immunostaining for SIX2 and Lysotracker red staining indicating dying cells in the cortical region of an E14.5 littermate control and TCre; *Fgf8^flox/null^* mutant kidney. Asterisk in D indicates cell death in renal vesicle. **G)** Quantification of the percentage of SIX2 positive cells that are also lysotracker positive. Control (TCre; *Fgf8^flox/WT^*) n = 6, TCre; *Fgf8^flox/null^* n = 4. Error bars represent +/- SEM, Student’s T-test was used to determine significance: *p < 0.05.

### Genetic abrogation of cell death does not adversely affect kidney development

One hypothesis for why the kidney fails to develop beyond a rudimentary state in the *Fgf8* mutant is that the NPs and their derivatives are dying, thereby preempting nephron formation. To explore this hypothesis, we attempted to restore cell survival by inactivating the redundantly required pro-apoptotic pathway genes *Bak* and *Bax* [18–20]. Homozygous null *Bak* mutants are normal and viable, but a majority of the homozygous *Bax* mutants die perinatally [11]. Combined loss of *Bak* and *Bax* resulted in defects in the patterning of certain tissues, such as interdigital webbing and imperforate vagina; however, the kidneys were reported to be overtly normal [11]. Therefore, we utilized *Bak* and *Bax* null alleles in the cross illustrated in Figure 2A to produce the three resultant genotypes of interest at the expected frequency of 1 in 8. The “most wildtype” genotype produced from this cross, hereafter called “*Control Bak*”, is lacking one allele of *Fgf8* in the TCre lineage and is homozygous null for *Bak*. The simplest genotype lacking Fgf8 produced from this cross, hereafter called *“Fgf8 mutant”*, lacks both alleles of *Fgf8* in the TCre lineage and is homozygous null for *Bak*. Lastly, we designated the genotype lacking *Fgf8* in the TCre lineage and also homozygous null for both Bak and Bax, “*Rescued Fgf8 mutant*” (Figure 2A).

**Figure 2.**
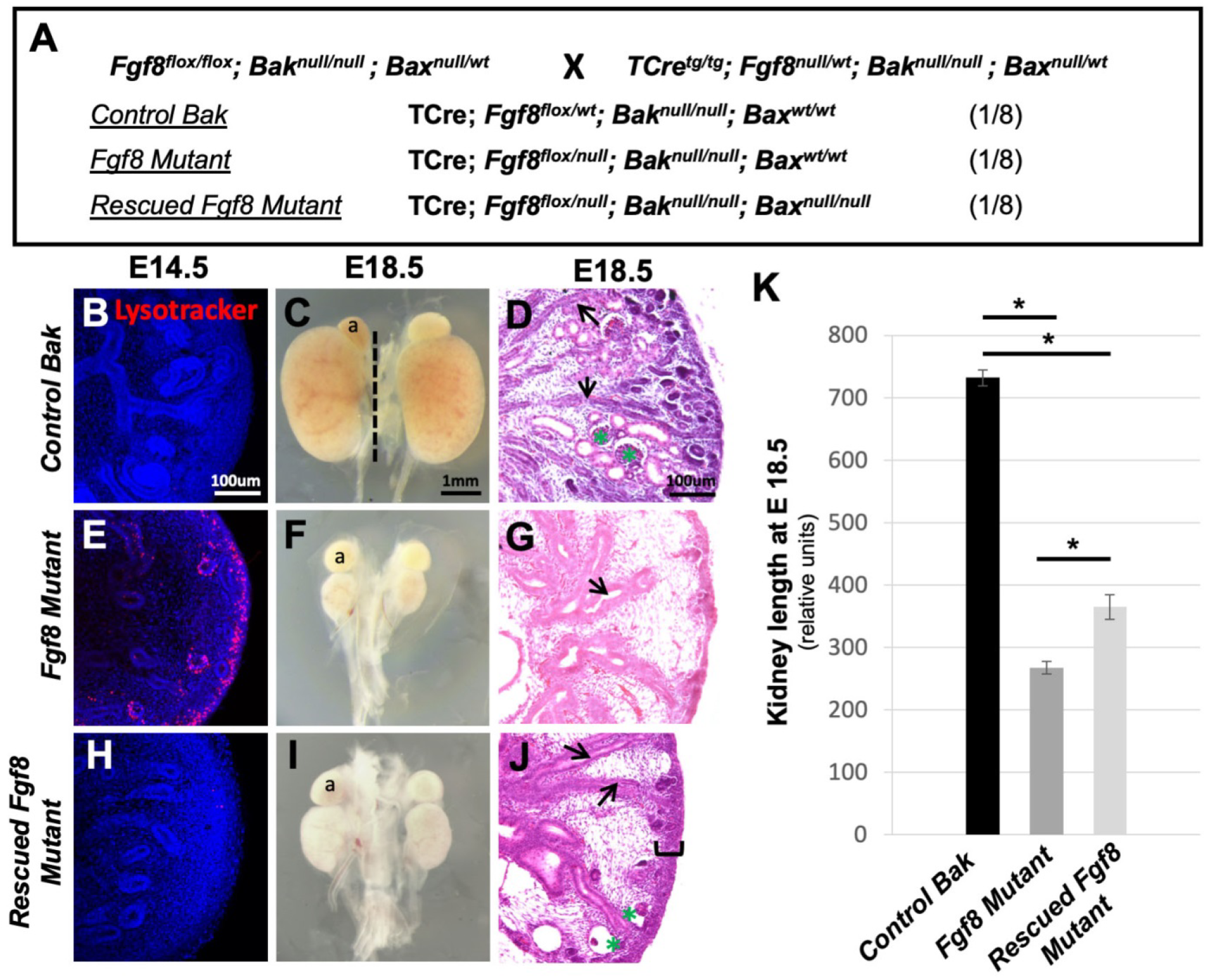
Loss of Bak and Bax rescues aberrant cell death in *Fgf8* mutants. **A)** Genetic cross to produce the three genotypes of interest listed below with assigned shorthand names (underlined). Note each genotype occurs at a frequency of 1/8; embryos heterozygous for Bax were not analyzed. **B, E, H)** Mid-sagittal sections of lysotracker stained E14.5 kidneys showing cell death; sections are counterstained with DAPI. **C, F, I)** Bright field image of whole E18.5 kidneys. Note, comparative size of adrenal gland (a) and kidney. **D, G, J**) Mid-sagittal thin sections of E18.5 kidneys. Black arrows point to tubules of uretic bud and green asterisks denote glomeruli. **K)** Graph of measurements of the length of E18.5 kidneys. Data represent the average length of the right and left kidney, dotted line in C indicates example of measurement. Error bars represent +/- SEM, Student’s t-test was used to determine significance: *p < 0.05. n = a minimum of 3 for all images. For K, *Control Bak* n = 6, *Fgf8 Mutant* n = 12, *Rescued Fgf8 Mutant* n = 5.

To determine if loss of *Bak* has any consequence on kidney development, we analyzed the morphology and size of wild-type *Bak* kidneys in comparison with *“Control Bak”* kidneys and found no changes (Supplemental Fig 1A-J, U), validating the use of the *Control Bak* kidneys as littermate controls that have normal kidney development. Likewise, the littermate *Fgf8 mutant* genotype was also homozygous null for *Bak*; however, we also found no discernible changes in cell death, morphology, or size of the kidneys when compared to TCre; *Fgf8^flox/null^*. mutants (Supplemental Fig 1K-U). We therefore used these *Fgf8 mutant* kidneys for comparison as littermate controls with the *Fgf8* loss-of-function kidney defect for the remainder of this study.

We then asked whether homozygous loss of both *Bak* and *Bax* impacts kidney development independent of *Fgf8*. Therefore, we scrutinized embryonic kidneys from animals containing one wild-type allele of *Fgf8* in the TCre lineage and null for both *Bak* and *Bax* and compared them with littermate *Control Bak* kidneys (therefore the only genetic difference is whether they are wild-type or null homozygous for *Bax*). These Control kidneys lacking both *Bak* and *Bax* were overtly normal by gross examination as previously noted (Supplemental Fig 2A, B)[11]. However, we noted they were 9.3% smaller in length than *Control Bak* kidneys (Supplemental Fig 2C). This change in length does not impact the number of nephrons on the basis of glomeruli counts or histological appearance of the kidney (Supplemental Fig 2D-F). Therefore we concluded this small size reduction did not diminish the utility of deleting *Bak* and *Bax* to interrogate the role of cell death in the phenotype caused by *Fgf8* loss in the kidney.

### Restored cell survival in *Fgf8 mutants* partially rescues kidney development

Having thoroughly characterized kidney development in our control littermate genotypes, we then examined the outcome of homozygous deletion of *Bak* and *Bax* to rescue the aberrant cell death in the *Fgf8 mutant* cortical regions. *Fgf8 mutants* have extensive cell death in the cortical regions and renal vesicles at E14.5 (Figure 2E compare to B), whereas *Rescued Fgf8 mutants* have no cell death in the cortical nephrogenic zone (Figure 2H). Compared to *Fgf8 mutants, Rescued Fgf8 mutant* kidneys exhibit an increase in size at E18.5 as measured by length (Figure 2I compared to 2K, quantified in 2F). However, these *Rescued Fgf8 mutant* kidneys are still significantly smaller than *Control Bak* kidneys (Figure 2C and 2K). *Rescued Fgf8 mutant* kidneys also displayed two histological changes: 1) an accumulation of blastemal-like mesenchymal cells in the cortical region (Figure 2J, black bracket) and 2) patent glomeruli (Figure 2J, green asterisks). Both observations are analyzed in greater detail in the following sections.

### Restoring Survival Rescues Nephron Progenitor Loss

To explore the apparent thickening of the cortical mesenchyme, we first stained kidneys with an antibody for SIX2 to detect nephron progenitors in the cap mesenchyme at E14.5, E16.5, and E18.5. *Fgf8 mutants* exhibited a normal pattern of SIX2 expression at E14.5. However by E16.5, NPs were markedly diminished, and by E18.5, almost no SIX2 positive cells were evident (Figure 3D-F). *Rescued Fgf8 Mutants* however had normal expression of SIX2 at E14.5 and E16.5 and at E18.5 had a large number of SIX2-positive cells in the cortical mesenchyme (Figure 3G-I), although the organization of SIX2-positive NPs was aberrant, i.e., they were less compact and individual cells were more diffusely scattered at E18.5 (compare Figure 3I with 3C).

**Figure 3.**
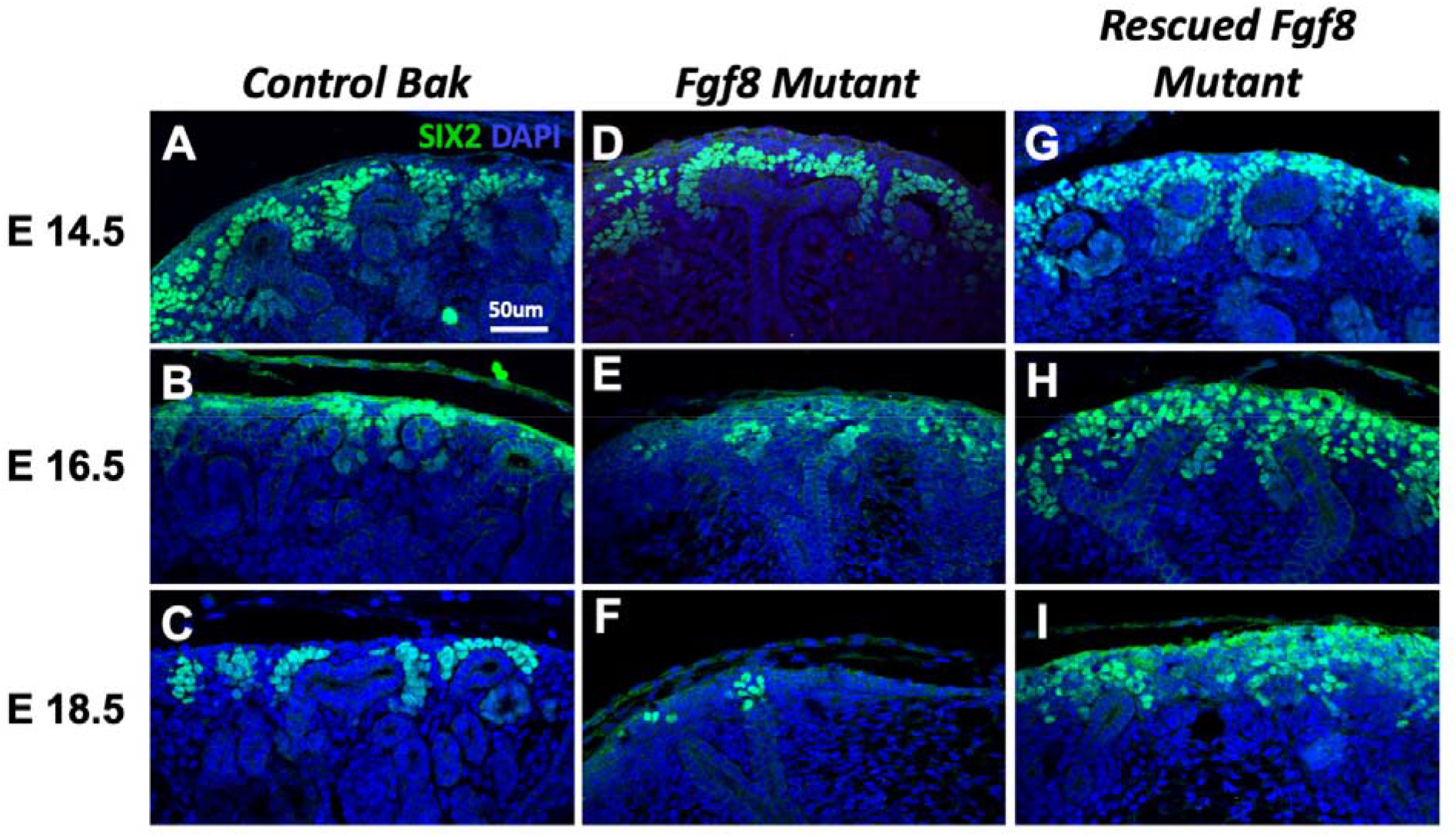
Restoring cell survival rescues loss of the Six2 progenitor population. **A-I)** Max intensity projections of mid-sagittal sections of SIX2 stained kidneys of noted genotypes at noted stages; sections are counterstained with DAPI. n = a minimum of 3 for all images.

To further examine the restoration of the NPs, we performed immunohistochemistry for a panel of MM markers. CITED1 labels a subset of the SIX2-positive NP population and is downregulated as these cells begin an mesenchymal-epithelial transition (Figure 4B) [21]. PAX2 is robustly expressed in the cap mesenchyme and renal vesicle and to a lesser extent the ureteric bud (UB) (Figure 4C) [22]. SALL1 is also highly expressed in cap mesenchyme and renal vesicle and at lower levels in stroma and UB (Figure 4D) [23]. We saw a similar trend in loss of MM markers as previously described [6]; SIX2, CITED1, PAX2, and SALL1 expression is substantially reduced or lost in *Fgf8 mutants* (Figure 4E-H). Furthermore, all of these markers are restored in the MM of *Rescued Fgf8 mutants* (Figure 4K-N). These observations were confirmed by RT-qPCR analyses, in which the levels of *Six2, Cited1*, and *Sall1* expression were significantly increased in *Rescued Fgf8 mutants* compared to *Fgf8 mutants* (Figure 4M). *Pax2* expression was restored to normal levels in *Rescued Fgf8 mutants*. This change, however, was found not to be significant when compared with expression in *Fgf8* mutants (Figure 4M). *Foxd1*, a marker of the stromal progenitor population, was unchanged among all genotypes (Figure 4M) [24]. Therefore, the restoration of cell survival in kidneys lacking *Fgf8* is sufficient to restore the MM progenitor populations that sustain nephrogenesis.

**Figure 4.**
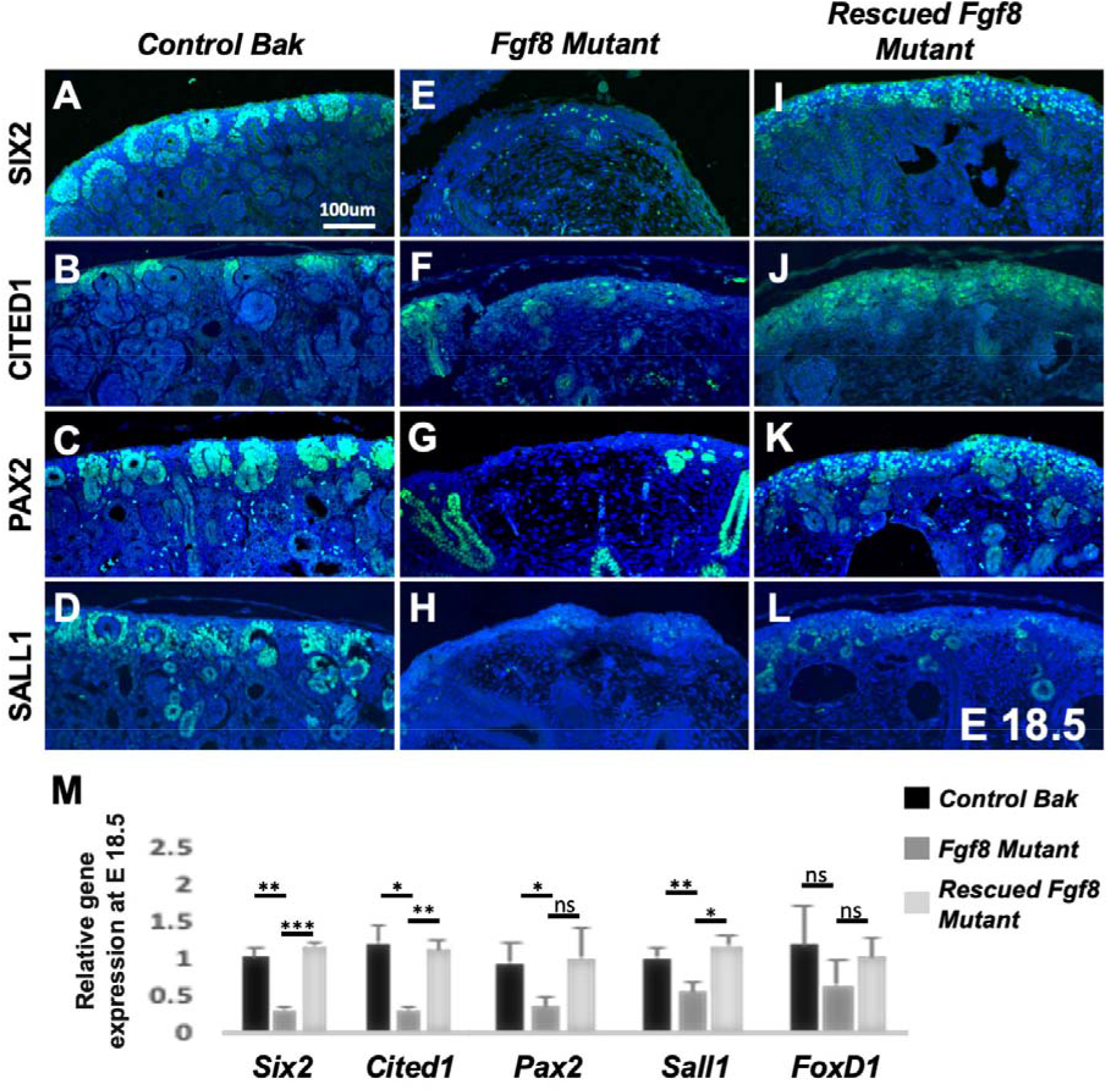
Restoring cell survival in *Fgf8* mutants rescues nephron progenitors. **A-L)** Max intensity projections of mid-sagittal sections of immunostained embryos for noted markers and genotypes at E18.5. n = a minimum of 3 for all images. **M)** Relative amount of gene expression of noted markers determined by quantitative PCR at E18.5. Error bars represent +/- SEM, Student’s t-test was used to determine significance: *p < 0.05, **p < 0.01, ***p < 0.001. n = 3 for genotypes and markers except *Control Bak* analysis for *Pax2* for which n = 6.

### Restoring Survival Does Not Rescue Nephron Formation

Histologically, we observed glomerular-like structures in *Rescued Fgf8 mutants* (Figure 2J). To confirm their identity, we immunolabeled E18.5 kidneys with an antibody for NEPHRIN, a marker of podocytes within glomeruli [25]. *Fgf8 mutant* kidneys rarely revealed any NEPHRIN-positive glomeruli (Figure 5B); however, *Rescued Fgf8 mutants* contained some NEPHRIN-positive glomeruli (Figure 5C), although fewer than *Control Bak* kidneys (Figure 5A). Quantification of NEPHRIN-positive glomeruli showed a significant increase in *Rescued Fgf8 mutant* kidneys (7.8 glomeruli per section) compared to *Fgf8 mutant* kidneys (1.45 glomeruli per section); however, *control Bak* kidneys had nearly 4-fold more glomeruli (29.2 glomeruli per section). Therefore, restoration of cell survival had a marginal, albeit statistically significant, effect on glomerular number.

**Figure 5.**
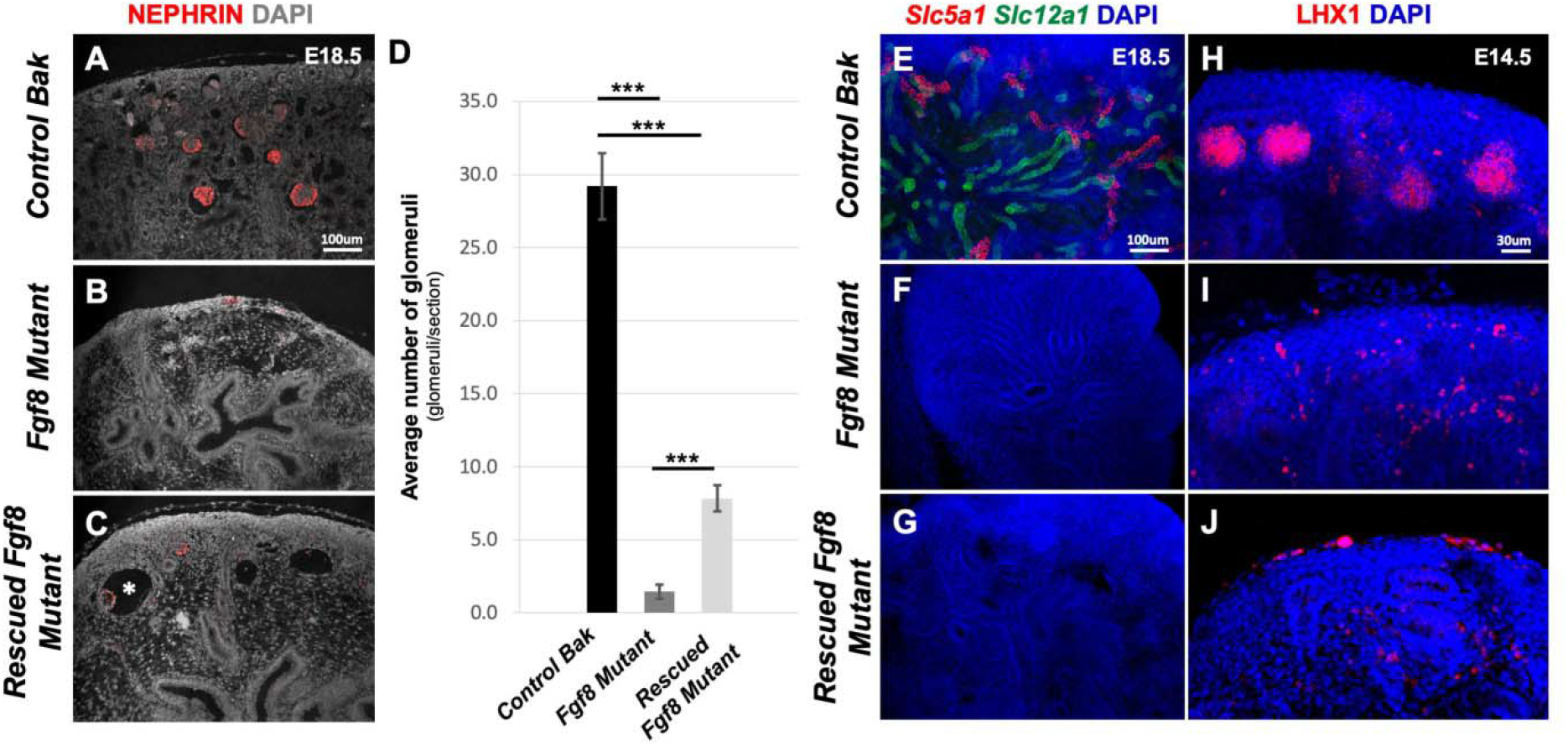
Restoring cell survival in *Fgf8* does not rescue nephrogenesis. **A-C)** Max intensity projections of mid-sagittal sections of NEPHRIN immunostained kidneys at E18.5. White asterisk in C notes cystic glomerulus. **D)** Quantification of glomeruli counts Error bars represent +/- SEM, Student’s t-test was used to determine significance: ***p < 0.001. *Control Bak* n = 5, *Fgf8 Mutant* n = 6, *Rescued Fgf8 Mutant* n = 6. **E-G)** Max intensity projections of mid-sagittal sections of *in situ* hybridization chain reaction (HCR) detection of *Slc5a1* and *Slcl2a1* mRNA at E18.5. **H-J)** Max intensity projections of mid-sagittal sections of LHX1 immunostained kidneys at E14.5. n = a minimum of 3 for all images.

The glomeruli of *Rescued Fgf8 mutants* appeared large and cystic (Figure 5C, asterisk), suggesting that these glomeruli lacked a patent tubular connection for drainage and waste excretion. We therefore examined whether there was any rescue of tubule formation when cell survival was restored in *Fgf8 mutants*. To accomplish this, we utilized fluorescent whole mount *in situ* hybridization chain reaction (HCR) using probes against *Slc5a1*, a marker of the proximal convoluted tubule [26], and *Slc12a1*, a marker of the ascending limb of the loop of Henle [27, 28]. *Control Bak* kidneys had clear labeling of *Slc5a1* and *Slac12a1* in each structure (Figure 5E), and, as expected, *Fgf8 mutants* had neither structure (Figure 5F). *Rescued Fgf8 mutants* also yielded no labeling from either marker, indicating that they too lack these renal tubules and thus functional nephrons (Figure 5G). Therefore, restoring cell survival in *Fgf8 mutants* did not rescue tubule formation.

LHX1 is a LIM homeobox transcription factor that is required for nephron formation, i.e., specifically tubule formation, and is expressed in pretubular aggregates, renal vesicles, and to a lesser extent, the UB [7]. We have previously shown that *Fgf8 mutants* lack *Lhx1* expression in pretubular aggregates and that *Lhx1* expression can be induced by FGF8 *ex vivo* [6]. We therefore examined LHX1 expression in mutants when cell survival is restored. As expected, LHX1 expression in the pretubular aggregates was present in littermate *Control Bak* kidneys displayed and absent in *Fgf8 mutants* kidneys (Figure 5H and I). *Rescued Fgf8 mutants* also had no pretubular aggregate expression of LHX1 (Figure 5J). Given the crucial role of LHX1 in tubulogenesis, this finding is consistent with the lack of tubules in *Rescued Fgf8 mutants* (Figure 5G).

In conclusion, blocking apoptotic cell death in *Fgf8* mutant kidneys can partially rescue kidney development, in which NPs are sustained, more glomerular-like structures appear, but subsequent nephron formation fails. These observations show that indeed FGF8 signals to block apoptosis in the cortical mesenchyme but also functions to promote subsequent nephronic development. It is possible that this latter signaling is necessary to induce *Lhx1* expression in the early stages of renal vesicle formation.

## Discussion

Here we analyze embryonic development in mouse kidneys that lack *Fgf8* in which we have restored cell survival through the genetic deletion of *Bak* and *Bax*. Our analysis is founded on the careful examination of control littermate kidneys; *Bak* and *Bax* deletion (in *Fgf8* wildtype kidneys) results in no overt consequence on renal development except for a slight decrease in kidney size. Our data show that FGF8 has potentially three functions during kidney development: 1) prevent nephron progenitor apoptosis 2) maintain *Lhx1* expression 3) drive NP cell migration. We consider each role below.

### FGF8 prevents nephron progenitor apoptosis

FGFs have been shown to promote cell survival in the developing lens, teeth, cranial neural crest, tail bud, and limbs [2, 29–32]. The aberrant cell death in developing kidneys, caused by *Fgf8* loss, is rescued when the intrinsic apoptotic pathway is abrogated, demonstrating that FGF8 maintains cell survival in a BAK/BAX*-*dependent manner. BAK and BAX act near the end of a cascade of activation events to permeabilize the mitochondrial outer membrane and release apoptotic factors [18, 20]. FGFs signal through multiple pathways [33] that potentially regulate different nodes of this cell survival/death pathway. For example, FGF signals have been shown to engage the PI3K/AKT pathway to phosphorylate BAD, resulting in antiapoptotic activity upstream of BAK/BAX [34, 35]. Another candidate for regulation by FGF-signaling is BIM. Deletion of one *Bim* allele, which encodes an apoptosis initiator that directly activates BAK/BAX [36, 37], will partially rescue aberrant cell death that underlie craniofacial defects that are due to the loss of *Fgfr1* and *Fgfr2* [29]. Our model for genetically rescuing cell death in the *Fgf8* mutant kidney presents a future opportunity to explore the role of FGFs in pro-survival signaling *in vivo*.

### FGF8 maintains Lhx1 expression

In addition to extensive cortical cell death, kidneys lacking *Fgf8* also fail to express *Lhx1*[6]. Pretubular aggregates form in *Lhx1*-deficient embryos but stall at renal vesicle formation and subsequently degenerate [7]. In the *Rescued Fgf8 mutant*, nephron progenitors are present, however renal vesicle formation similarly arrests shortly after initiation. Restoration of cell survival does not restore LHX1 protein expression in *Fgf8 mutant* kidneys. Therefore, in addition to providing a pro-survival signal, FGF8 is likely also required for inducing or maintaining *Lhx1* expression. Deficient *Lhx1* expression could explain the lack of renal vesicle maturation and subsequent failure of nephron formation in the *Rescued Fgf8 mutant*. The renal vesicles that are found in the *Rescued Fgf8 mutant* at E18.5 are likely formed prior to E14.5 and would undergo apoptosis in the absence of *Fgf8*, if apoptosis were not blocked. However, due to deficient *Lhx1* expression, these primitive renal vesicles fail to elongate and form mature nephrons. It is possible that aberrant cell death is a function of *Lhx1* loss. Although apoptosis is yet to be analyzed in *Lhx2* mutant kidneys, like *Fgf8* mutant kidneys, they are smaller [7]. Moreover, *Lhx2* loss has been shown to cause increased apoptosis in the Mu□llerian duct epithelium [38].

The initial formation of renal vesicles prior to E14.5 in the Fgf8 mutant is likely driven by redundant signaling from FGF9 and FGF20. *Fgf9* is expressed in the UB and weakly in the MM, while *Fgf20* is found in the MM and early renal vesicle [39]. When *Fgf9* and *Fgf20* are lost, there is a rapid depletion of MM between E11.5 and E12.5 involving apoptosis of this tissue [39]. *Fgf9* and *Fgf20* are however not sufficient for sustaining renal vesicle development beyond E14.5, at which time *Fgf8* is required for this process. Therefore, these three FGF family members may act redundantly over the course of kidney formation.

### FGF8 drives NP cell migration

The phenotype of the metanephric mesenchyme in *Rescued Fgf8 mutants* suggests that *Fgf8* also functions in renal development by driving NP cell migration during renal vesicle formation. In the absence of *Fgf8*, renal progenitor cells, as labeled by SIX2, apparently fail to migrate and therefore accumulate in the cortical nephrogenic zone. Conditional inactivation of *Lhx1* in the metanephric mesenchyme results in cessation of renal vesicle development; however, nephron progenitors do not appear to accumulate in the cortex as they do in *Rescued Fgf8 mutants* [7]. We therefore speculate that this accumulation highlights a third role for *Fgf8* in kidney development, which is inducing migration of cortical NP cells at the initiation of renal vesicle formation.

Migration of NP cells is an essential process in both the maintenance of the cortical NPs as well as differentiation [40]. The cap mesenchyme (CM) cells exist in a niche surrounding the distal end of cortical uretic bud tips. Migration of CM is driven by a combination of attractive and repulsive cues from the surrounding interstitial cells and from uretic bud tips [41]. It is thought that cells must migrate out of this region to downregulate progenitor maintenance genes and begin initiating differentiation programs [40].

Starting at E11.5, *Fgf8* is expressed in a ring below the branching UB, and thereafter within the pretubular aggregates and newly formed renal vesicles at E14.5 NPs [6, 42] and therefore may encode as an attractant activity for cortical NPs and migrating migrating CM cells.

Therefore, it is consistent that a failure in migration of cortical nephron progenitor descendants would lead to an accumulation of SIX2-positive cells in the nephrogenic zone, as we observe in *Rescued Fgf8 mutants*. WNT signaling has been shown to be key in regulating CM migration, specifically WNT9b and WNT11 [43]. *Wnt11* inactivation causes a failure in CM migration and subsequent disorganization, similar to the *Rescued Fgf8 mutant* phenotype. FGFs and WNTs interact in multiple tissues to affect tissue morphogenesis [44–46], and it is therefore possible that the failed CM migration that occurs in the *Rescued Fgf8 mutant* is due to loss of WNT-driven migration.

FGF8 signaling within the kidney mesenchyme functions to support cell survival by blocking apoptosis, instructing cell fate, and possibly driving cell migration, all of which are related to basic hallmarks of cancer development and progression [47]. Furthermore, *Lhx1* and *Six2*, both regulated by FGF8, are implicated in renal cancer development [48, 49]. Further investigation of the mechanism of FGF8 signaling to control the expression of the genes and developmental processes will not only impact our understanding of the role of *Fgf8* in kidney development, but also lead to insights regarding cancer.

## Methods and Materials

### Alleles, Genotyping, and Kidney Dissection

All mice were kept on a mixed background. PCR-genotyping for each allele was performed using the following primer combinations, *Fgf8flox*[50] (5’- GG TCT TTC TTA GGG CTA TCC AAC and 5’- GCT CAC CTT GGC AAT TAG CTT C) *Fgf8Δ* [50] (5’-CCA GAG GTG GAG TCT CAG GTC C and 5’-GCA CAA CTA GAA GGC AGC TCC C), *Bak*[11] (5’- GGC TCT TCA CCC CTT ACA TCA G, 5’- GTT TAG CGG GCC TGG CAA CG, and 5’- GCA GCG CAT CGC CTT CTA TC), *BaxΔ* [11] (5’-CAA CTC CTA CCG CAA GTC CTG G and 5’- GAA CCC TAG GAC CCC TCC G), *BaxWT* (5’-TGC CGA ACT GGG CAC TGT TG and 5’- GTC CTG GGG AAT GTG GAC TG), TCre[6] (5’- GGG ACC CAT TTT TCT CTT CC and 5’-CCA TGA GTG AAC GAA CCT GG). Embryos where dissected on the noted embryonic day in chilled PBS and kidneys where removed from the embryo and fixed in 4% paraformaldehyde at 4°C, rocking overnight. The following day, kidneys where washed in PBS, then dehydrated, stepwise, into 100% methanol and stored at −20 °C.

### Wholemount Lyostracker Staining

E14.5 and E16.5 kidneys were dissected in 37° C Hanks BSS without phenol red with Mg^+2^ and Ca^+2^. The isolated kidneys were placed in 5 ml Hanks and 25 ul LysoTracker Red (Invitrogen) and cultured at 37° C for 30 minutes, then rinsed in Hanks and fixed in 4% paraformaldehyde at room temperature for 30 minutes. They were then rinsed with PBS, dehydrated into 100% methanol and stored at −20° C, until time of imaging.

### Lysotracker Staining and Six2 Immunohistochemistry

Lysotracker staining was performed as noted above. Then kidneys were rehydrated and rinsed in 1% Triton in PBS for 10 minutes at room temperature. The tissues were then blocked in 5% goat serum in 1% Triton X-100 PBS for two hours at room temperature and incubated in anti-SIX2 (Proteintech: 11562-1-AP, 1:50) overnight at 4°C while rocking. The following day, kidneys were then washed in 1% Triton X-100 in PBS 3 x 10 minutes at room temperature and incubated with Alexa Fluor 488 conjugated goat anti-rabbit IgG (Thermo Fisher: A-11008, 1:250) and 0.5ug/mL DAPI overnight at 4°C. Kidneys were then washed with 1% Triton X-100 in PBS 3 x 10 minutes at room temperature, embedded in 7% Low Melting Point Agarose, and 80 um sections were cut using a vibratome.

### Immunohistochemistry on Paraffin Sections

Kidneys were moved into 100% ethanol, then into Citrisolv, followed by two changes of melted paraffin wax, before final positioning in paraffin. Sections (8um) where cut from the paraffin blocks, and deparaffinized using Citrisolv reagent. Sections where blocked with 5% goat serum in 1% Triton X-100 PBS for two hours at room temperature and incubated in primary antibody overnight at 4°C in a humidified chamber. Slides where then washed 3x 20 minutes in 1% Triton X-100 PBS at room temperature and incubated with the appropriate secondary antibody and 0.5ug/mL DAPI overnight at 4°C in a humidified chamber. Slides were washed 3x 20 minutes in 1% Triton X-100 PBS at room temperature and tissues preserved under coverslips using Aqua-Poly/Mount (Polysciences, 18606). The following primary and secondary antibodies at indicated concentrations were used: rabbit polyclonal Six2 antibody (Proteintech, 1:50), rabbit polyclonal Cited1 antibody (Neomarkers, 1:50), rabbit polyclonal Pax2 antibody (Invitrogen antibodies, 1:50), rabbit polyclonal Sall1 antibodies (abcam, 1:50), and guinea pig polyclonal Nephrin antibodies (Progen, 1:50), Alexa Fluor 488 conjugated goat anti-rabbit IgG (Thermo Fisher: A-11008, 1:250), Alexa Fluor 647 conjugated goat anti-guinea pig IgG (Thermo Fisher: A-21450, 1:250).

### Confocal Imaging and Image Analysis

All fluorescent images were taken using an Olympus FV1000 with a 20x UPlanApo objective (NA: 0.70) with 3X-Kalman averaging. Images where processed using FIJI software to generate Max Intensity Projections. Identical intensity ranges where used between compared samples.

### Determination of SIX2-Lysotracker double-positive cells

Spatial models were generated using Imaris software (Imaris V9.2.1, Bitplane Inc), to analyze Lysotracker-SIX2 double positive cells. This was done to account for nuclear-localized SIX2 staining and lysotracker staining, which may be localized anywhere within the cell, since standard pixel-based colocalization cannot accurately identify double positive cells. Within the Imaris software, the “spot” modeling tool was used to generate spot features for each channel (SIX2 and Lysotracker). Because the diameter and intensity of SIX2 positive cells varied, three different spot models were created: 7 um XY diameter and 7 um Z diameter using a minimum sum intensity threshold of 1.25 e5; 2 um XY diameter and 4 um Z diameter using a minimum sum intensity threshold of 5800 and upper threshold of 1.50 e4; and 3 um XY diameter and 4 um Z diameter using a minimum sum intensity threshold of 8000 and upper threshold of 1.50 e5. Redundant spots generated from the three models that labeled the same SIX2 positive cells were eliminated from the analysis by colocalizing spots that were within 5 um from each other, and only analyzing those that were unique (omitting the redundant spots from the analysis). Each SIX2 spot labeled one nucleus and therefore one cell. For spot modeling of the Lysotracker positive signal a 3.5 um XY diameter and 8 um Z diameter and minimum sum intensity of 1.00 e4 was used to generate spots.

To exclusively analyze Lysotracker staining in the cortical region (and not renal vesicle or medullary cell death) the Imaris “surface” tool was utilized to generate a volumetric model. Surface models were generated based on SIX2 positivity with surface detail of 2 um and a low absolute intensity threshold of 1200, to encompass all SIX2 positive cells. All lysotracker spots within 10 um of the Six2 positive surface model were considered cortical and used for colocalization with spots modeling SIX2 positive cells. The threshold for colocalization between SIX2 and lysotracker spots was 5 um. Colocalization of one SIX2 spot with multiple lysotracker spots or vice versa was manually checked to ensure single colocalized pairs; in the event of multiple colocalizations, only one of each class of spot was counted to be colocalized.

### Hybridization Chain Reaction whole mount *in situ*

Hybridization Chain Reaction whole mount *in situ* split initiator probes (V3.0) were designed and synthesized by Molecular Instruments, Inc. Staining was carried out as previously described [51].

### RT-qPCR

Both kidney rudiments from each embryo were isolated at the indicated ages, frozen on dry ice, and stored at −80°C, while the embryos were being genotyped. Total RNA from 10-15 pairs of metanephroi with the same genotypes was purified using TRIzol reagent (Invitrogen cat# 15596026). Following extraction, RNA preparations were dissolved in 10 μl DEPC-treated distilled water. For RT-PCR, random hexamer primers were used to generate cDNA with a Verso kit from Thermo Scientific cat #AB1453/B. For this, total RNA was incubated at 42°C for 30 min and 95°C for 2 min. For quantitative PCR, 1 μl cDNA was used in a 20 μl reaction mixture with a Bio-Rad SsoFast EvaGreen super mix Kit (cat# 1725201) and 3μM gene-specific primers in a Bio Rad CFX96 real-time PCR detection system. Incubation conditions typically consisted of 45 cycles (3min 95°c, 5s 95°c 5s 65°c), depending upon primer optimal annealing temperatures. Relative levels of gene expression were calculated using ΔCq values and normalized to *GAPDH* expression. Primers for each cDNA gene-specific reaction were as follows: *Cited1* (5’ - AAT GTG TCC GTC GTG GAT CTG - 3’ and 5’ - CTG CTT CAC CAC CTT CTT GAT GT - 3’); *FoxD1* (5’ – TCG CTC TGT CTT GGC ACT AGG A - 3’ and 5’ – ACG CCT GGA CCT GAG AAT CTC TAC - 3’);; *GAPDH* (5’ - AAT GTG TCC GTC GTG GAT CTG - 3’ and 5’ - CTG CTT CAC CAC CTT CTT GAT GT - 3’); *Pax2* (5’- AGG CAT CAG AGC ACA TCA AAT CAG - 3’ and 5’ - GGG TTG GCC GAT GCA GAT AG - 3’); *Sall1* (5’ - TGT CAA GTT CCC AGA AAT GTT CCA and 5’ - ATG CCG CCG TTC TGA ATG A - 3’); and *Six2* (5’ - GCA ACT TCC GCG AGC TCT AC - 3’ and 5’ - GCC TTG AGC CAC AAC TGC TG - 3’).

---

**Supplementary Figure 1.**
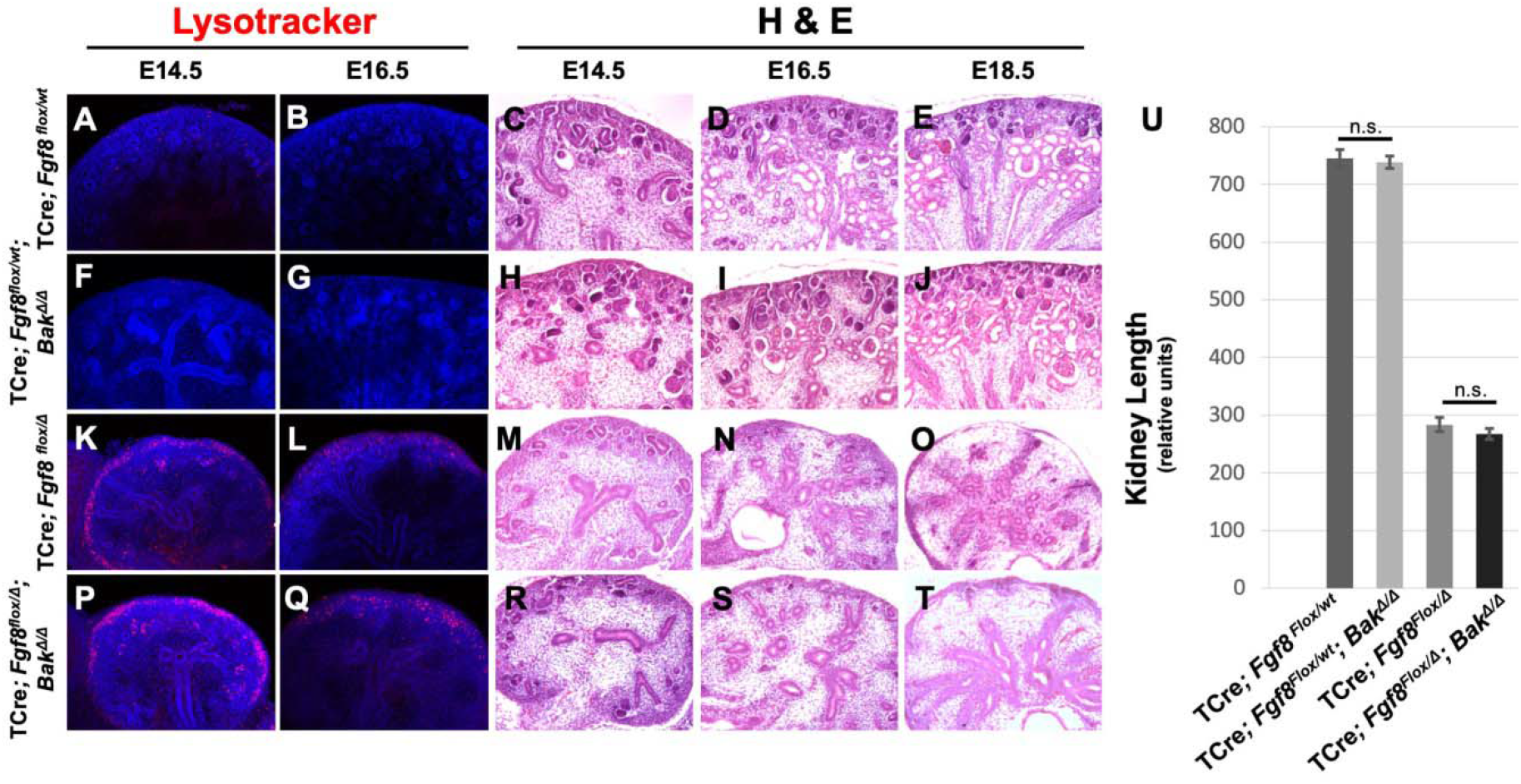
Loss of *Bak* has no patent phenotypic consequence on TCre; *Fgf8^flox/null^* mutants and littermate controls (TCre; *Fgf8^flox/WT^*). A, B, F, G, K, L, P, Q) Max intensity projections of mid-sagittal sections of lysotracker-stained kidneys at noted stages, counterstained with DAPI. **C-E, H-J, M-O, R-T)** H & E stained thin mid-sagittal sections at noted stages. **U)** Quantification of kidney length in relative units. Error bars represent +/- SEM, Student’s t-test was used to determine significance. TCre*; Fgf8 ^Flox/wt^* n = 13, TCre*; Fgf8^Flox/wt^; Bak^Δ/Δ^* n = 4, TCre*; Fgf8^Flox/Δ^* n = 9, TCre; *Fgf8^Flox/Δ^; Bak^Δ/Δ^* n = 9.

**Supplementary Figure 2.**
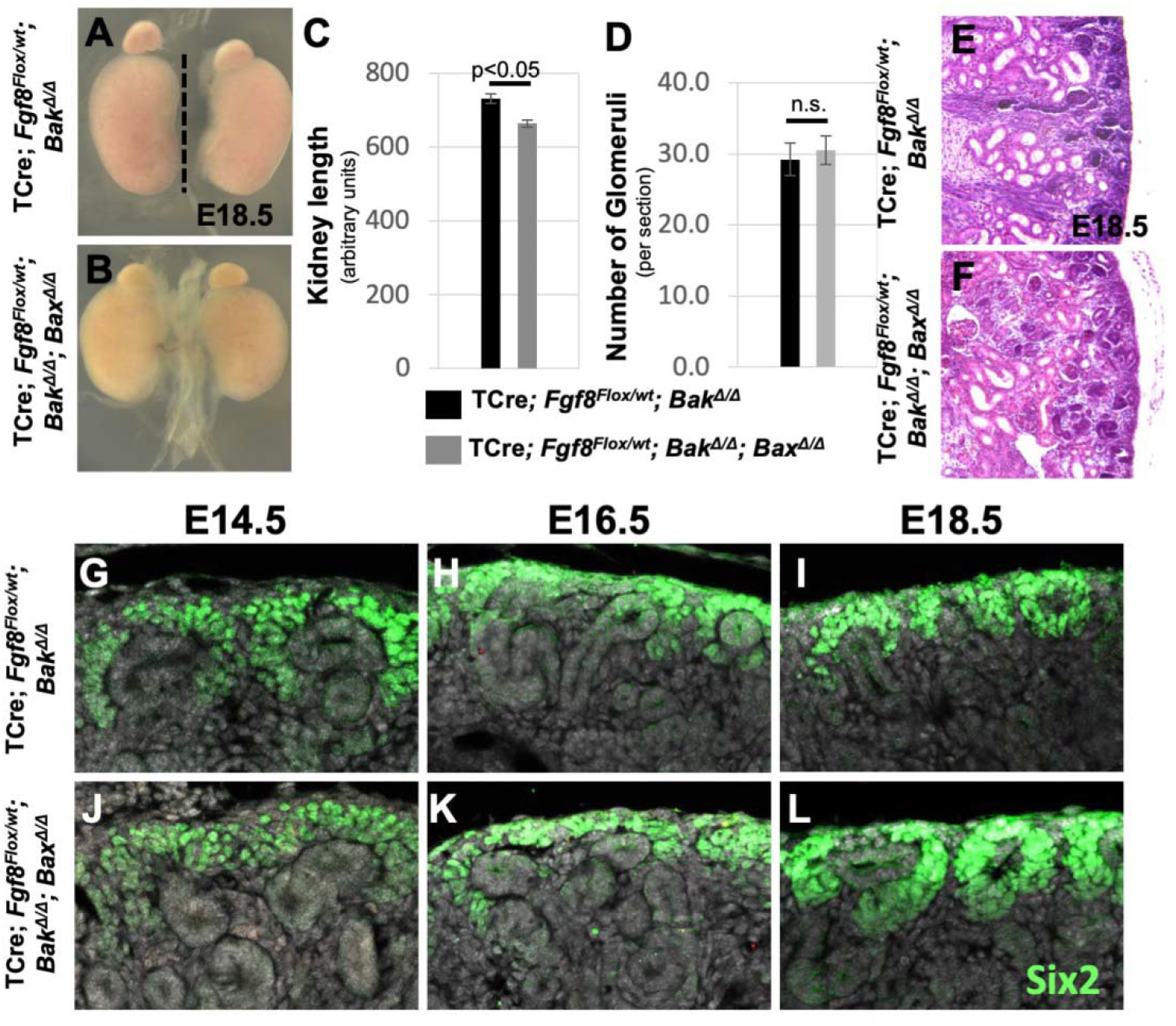
Inactivation of Bak and Bax has minimal effects on kidney development. **A, B)** Brightfield images of kidneys of noted genotypes at E18.5. **C)** Anterior-posterior length of kidneys of noted genotypes at E18.5; dotted line in A denotes measurement taken. **D)** Average number of glomeruli per mid-sagittal section of noted genotypes; glomeruli counted from E, F. **E, F)** H & E stained mid-sagittal sections of E18.5 kidneys of noted genotypes. G-L) Max intensity projections of SIX2 and DAPI stained regions of noted genotypes at E14.5, E16.5, and E18.5. C, D, error bars represent SEM, t-test was used to determine significance.

## Acknowledgements

We thank Patricia Abete-Luzi, Christian Bonatto, Michael Boylan, Cindy Elder, and Erika Truffer for technical assistance or critical reading of the manuscript.

